# Cellular COPI components promote geminivirus infections by facilitating the chloroplast localization of viral C4/AC4 proteins

**DOI:** 10.1101/2023.11.08.566295

**Authors:** Wenhao Zhao, Yinghua Ji, Yijun Zhou, Xiaofeng Wang

## Abstract

Geminiviruses are a family of viruses that infect numerous crops and cause extensive agricultural losses worldwide. During viral infection, geminiviral C4/AC4 proteins relocate from the plasma membrane (PM) to chloroplasts, where they inhibit chloroplast-mediated host defense, including the biosynthesis of salicylic acid (SA). However, how are C4/AC4 proteins transported to chloroplasts is unknown. We report here that the Coat Protein I (COPI) components play a critical role in redistributing Tomato yellow leaf curl virus (TYLCV) C4 protein to chloroplasts. TYLCV C4 interacts with the β subunit of COPI, and the coexpression of both in *Nicotiana benthamiana* cells promotes the enrichment of C4 in chloroplasts, which also occurs during TYLCV infection and is blocked by an inhibitor of the COPI pathway. Overexpression of COPI components promotes but knockdown of gene expression inhibits TYLCV infection. The COPI pathway plays similar roles in C4/AC4 transport and infections of other geminiviruses, including Beet curly top virus and East African cassava mosaic virus. Our results identify an unconventional role of the COPI pathway in protein trafficking to chloroplasts during geminiviruses infections in plants, and suggest a broad-spectrum antiviral strategy in controlling geminiviruses by manipulating COPI components.

## Introduction

Geminiviruses are a class of single-strand DNA viruses that cause tremendous agricultural losses worldwide (Fondong et al., 2007; Padmanabhan et al., 2022). Tomato yellow leaf curl virus (TYLCV) is a representative member of the *Geminiviridae* family and a monopartite begomovirus that replicates in the nucleus. The C4 protein of TYLCV plays multiple roles during viral infection. C4 localizes to the plasmodesmata (PD) where it strongly associates with the PD-localized receptor-like kinase proteins (RLKs) and suppresses the cell-to-cell spread of RNA interference signals moving through the PD (Rosas-Diaz et al., 2018). C4 also inhibits chloroplast-dependent SA biosynthesis to promote viral infection (Medina-Puche et al., 2020).

C4 and AC4 proteins from different geminiviruses, such as TYLCV, Beet curly top virus (BCTV), and East African cassava mosaic virus (EACMV), primarily localize to the plasma membrane (PM) and weakly to chloroplasts when expressed in plant cells (Medina-Puche et al., 2020). These proteins have two overlapping localization signals: an N-myristoylation site (Glycine2) that is critical for the PM localization, and a chloroplast transit peptide (cTP) that targets chloroplasts (Medina-Puche et al., 2020; Rosas-Diaz et al., 2018). Interestingly, geminivirus infections or the expression of viral Rep protein redirects a large fraction of PM-bound C4/AC4 protein to chloroplasts where C4 interacts with the thylakoid membrane-bound plant calcium-sensing receptor (CAS) and acts as a suppressor of SA biosynthesis (Rosas-Diaz et al., 2018; Krenz et al., 2010; Teixeira et al., 2021). An intriguing question is how these PM-localized C4/AC4 proteins are translocated to chloroplasts.

Chloroplasts are one of the most dynamic and important organelles in plant cells with a major function for photosynthesis. Chloroplasts are also one of the major sites for the synthesis of fatty acids, membrane lipids, amino acids, and starch (Chan et al., 2016). Additionally, phytohormones involved in defense responses and inter-organelle signaling are produced in chloroplasts, including reactive oxygen species (ROS), salicylic acid (SA), and jasmonic acid (JA) precursors, and therefore, chloroplasts play active roles in mitigating biotic and abiotic stresses (Gan et al., 2019; Caplan et al., 2008; de Torres Zabala et al., 2015; Sowden et al., 2017; Medina-Puche et al., 2020; Wang et al., 2021). During host immune responses to pathogen infections, multiple chloroplasts align around the nucleus and establish physical contact with the nucleus via stromules, which are tubular membrane projections of the chloroplasts (Kumar et al., 2018; Kwok and Hanson, 2004). Plant immune response signals, such as H_2_O_2_ and SA, are transferred to the nucleus where these signals regulate expressions of defense genes or induce programmed cell death (Natesan et al., 2005; Caplan et al., 2015; Exposito-Rodriguez et al., 2017). Perinuclear clustering of chloroplasts was further shown to be a general host response to the perception of pathogen infections or expression of certain viral proteins (Ding et al., 2019).

Trafficking of proteins among organelles, such as the endoplasmic reticulum (ER), the Golgi apparatus, and the trans-Golgi network (TGN), and the membrane, is mediated by coat protein complex I (COPI) and II (COPII) pathways (Paul and Frigerio, 2007; Feng et al., 2016; Luo and Boyce, 2019; Wessels et al., 2006). COPI vesicles, which originate from Golgi membranes and are involved in intra-Golgi and Golgi to ER protein retrieval, consist of seven subunits, α, β, β’, δ, ε, γ, and ζ (Waters et al., 1991; Arakel and Schwappach, 2018; Bethune and Wieland, 2018). The initiation of the COPI assembly starts with a Golgi-localized GTPase Arf1 (ADP ribosylation factor 1), which cycles between GDP- and GTP-bound forms. A guanine nucleotide exchange factor, GBF1, activates Arf1 by changing GDP-bound to GTP-bound Arf1. Arf1-GTP subsequently recruits COPI components to assemble into coatomer complexes (Beck et al., 2009). COPI/II pathways were also shown to be involved in viral proteins trafficking during viral infections (Richardson et al., 2014; Barajas and Nagy, 2010; Cabanillas et al., 2018; Agaoua et al., 2021; Solovyev et al., 2022; Sangeetha and Jebasingh, 2021).

In this study, we demonstrate a proviral role of tomato (*Solanum Lycopersicum*) COPIβ (SlCOPIβ) in TYLCV C4 trafficking and viral infection. SlCOPIβ specifically interacts and colocalizes with TYLCV C4 in chloroplasts. Knocking down the expression of *SlCOPIβ* inhibits and overexpressing *SlCOPIβ* promotes TYLCV infection in tomato plants. We further show that coexpression of C4 and SlCOPIβ promotes the clustering of chloroplasts along the nucleus, a phenomenon that occurs during TYLCV infection, expression of TYLCV Rep protein, or upon treatment of the epitope peptide of bacterial flagellin, flg22 (Ding et al., 2019). We additionally demonstrate that the COPI pathway is required for targeting C4 to chloroplasts and for the perinuclear clustering of chloroplasts. More importantly, SlCOPIβ interacts with and redistributes BCTV C4 and EACMV AC4 to chloroplasts, and is positively involved in BCTV infection, suggesting a proviral role of SlCOPIβ in geminivirus infections. Our work demonstrates a novel role of the COPI pathway in promoting geminiviral infections by translocating viral C4/AC4 proteins to chloroplasts, and suggests a new way for viral protein trafficking and viral infection.

## Results

### The TYLCV C4 protein interacts with tomato COPIβ

It has been reported that the TYLCV C4 protein localizes at the PM, PD, and chloroplasts (Rosas-Diaz et al., 2018). Our results, using a yellow fluorescent protein (YFP)-tagged C4, agreed with previous reports (Medina-Puche et al., 2020; Rosas-Diaz et al., 2018) that strong fluorescence signals were found at the PM and a minor fraction was detected in chloroplasts (Figure 1A and Supplemental Figure S1A). Additionally, coexpression of C4-YFP with a red fluorescent protein (RFP)-tagged Tobacco mosaic virus movement protein (TMV MP-RFP), which serves as a PD marker, confirmed that C4 localized to PD (Figure 1B).

**Figure 1.**
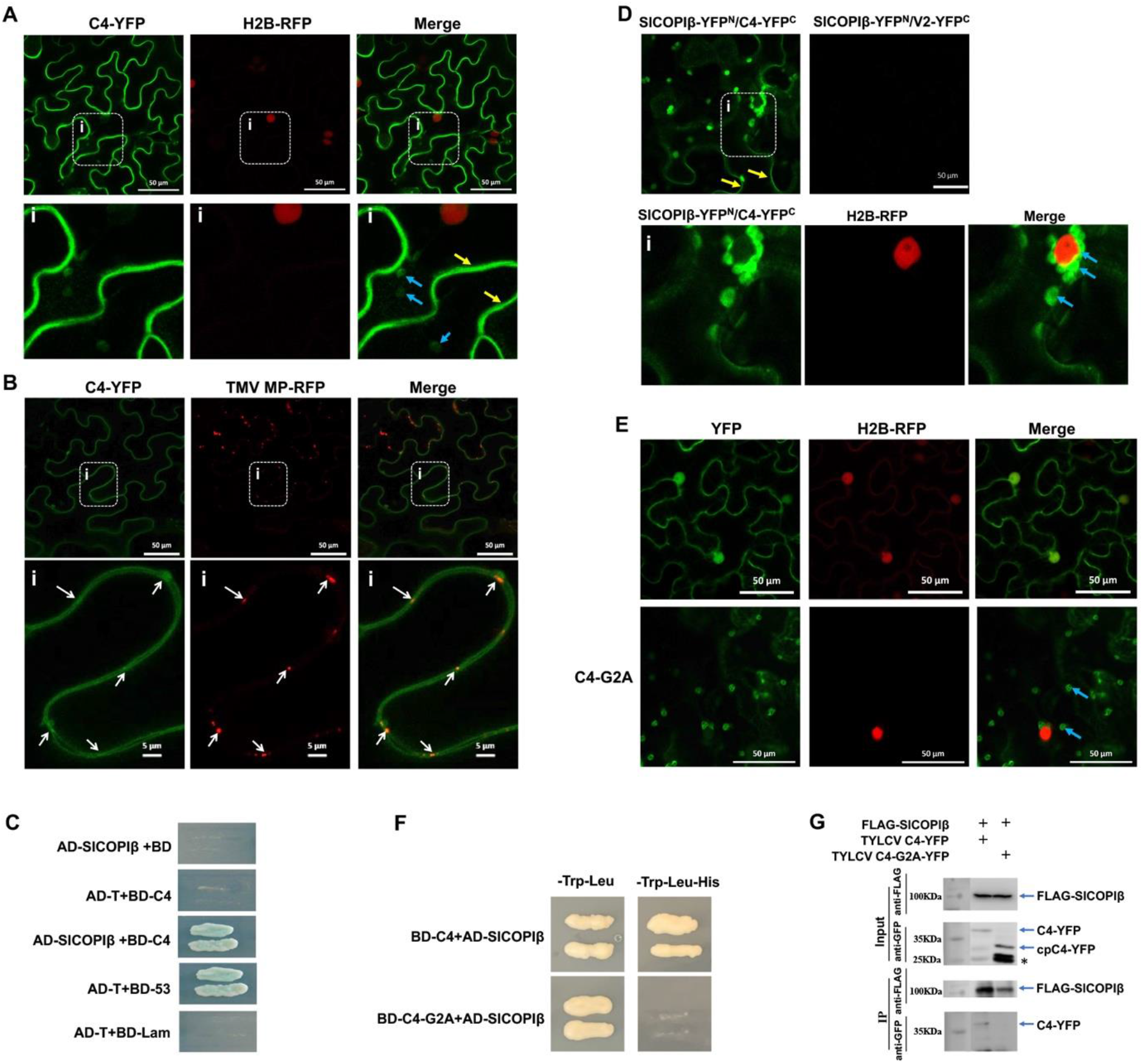
The specific interaction between TYLCV C4 and SlCOPIβ. (A) Subcellular localization of C4-YFP in RFP-tagged H2B transgenic *N. benthamiana* cells. Yellow arrows point to the PM; blue arrows point to chloroplasts. The H2B-RFP signal represents the nucleus. The rectangle indicates the enlarged region blow. Bars: 50 μm. (B) C4-YFP colocalized with an RFP-tagged plasmodesmata (PD) marker, Tobacco mosaic virus movement protein (TMV MP-RFP), in *N. benthamiana* cells. Bars: 50 μm. White arrows indicate the punctate signals of PD. Bars: 5 μm. (C) The interaction between C4 and SlCOPIβ as revealed by a Y2H assay. Yeast cells expressing the different combinations were grown on the SD/-His/-Leu/-Trp/-Ade selective medium with X-α-Gal. AD-T and BD-53 or AD-T and BD-Lam was used as a positive or negative control, respectively. (D) A BiFC assay to confirm the C4-SlCOPIβ interaction in *N. benthamiana* cells. Yellow arrows point to the PM; blue arrows point to the chloroplasts. Bars: 50 μm. (E) The localization of C4-G2A-YFP in H2B-RFP transgenic *N. benthamiana* cells. Blue arrows point to chloroplasts in cells. Bars: 50 μm. (F) SlCOPIβ does not interact with C4-G2A as revealed by a Y2H assay. (G) A co-immunoprecipitation (co-IP) assay to test the interaction between SlCOPIβ and C4 or C4-G2A in plant cells. Total leaf lysates coexpressing FLAG-SlCOPIβ and C4-YFP (Lane 1) or FLAG-SlCOPIβ and C4-G2A-YFP (Lane 2) were cleared and incubated with anti-FLAG beads. Proteins associated with beads were analyzed on a SDS-PAGE gel and immunoblotted using anti-GFP or -FLAG antibody. C4-YFP indicates the full-length, PM-localized C4-YFP and cpC4-YFP indicates the processed, chloroplast-localized form. Asterisks indicate unidentified bands and/or potential degradation products of C4-YFP.

Given the versatile functions of C4, we set to identify host proteins that interact with C4. We screened a yeast two-hybrid cDNA library, which was constructed by using mRNA isolated from tomato leaf tissues. A single cDNA encoding tomato COPIβ (SlCOPIβ) (GenBank accession no. XM_026027730) was identified. We next amplified full-length coding sequence of SlCOPIβ from tomato leaf tissue and confirmed the interaction in the Y2H assay (Figure 1C).

We next verified the C4-SlCOPIβ interaction in plants cells using a bimolecular fluorescence complementation (BiFC) assay. SlCOPIβ-YFP^N^ and C4-YFP^C^ were co-infiltrated into *N. benthamiana* leaves and strong fluorescence signals were observed in both the PM and chloroplasts (Figure 1D), reinforcing the C4-SlCOPIβ interaction in plant cells.

Localization of C4 protein at the PM/PD depends on the presence of a myristoylation motif at the N terminus and PM/PD localization of C4 protein is required for virus infectivity (Medina-Puche et al., 2020; Rosas-Diaz et al., 2018). A non-myristoylable mutant of C4 (C4-G2A) was fused to YFP and fluorescence signal was observed at 40 hours post agroinfiltration (hpai). In >90% of cells (30 cells), the C4-G2A-YFP fluorescence signal was predominantly found in chloroplasts but not in the PM/PD (Figure 1E and Supplemental Figure S1B), in agreement with what has been reported (Medina-Puche et al., 2020; Rosas-Diaz et al., 2018). We next tested whether C4-G2A interacted with SlCOPIβ using Y2H and a co-immunoprecipitation (co-IP) assay. As shown in Figure 1F, we found no interaction between SlCOPIβ and C4-G2A, in sharp contrast to that between SlCOPIβ and wild-type (WT) C4. In addition, when YFP-tagged WT C4 or C4-G2A were coexpressed with SlCOPIβ in *N. benthamiana* cells, only WT C4-YFP, but not C4-G2A-YFP, was pulled down along with FLAG-SlCOPIβ (Figure 1G). These results confirm that G2 is a key site in C4 for its interaction with SlCOPIβ.

### SlCOPIβ is a proviral factor in TYLCV infection in tomato plants

Because C4 was enriched at chloroplasts in the presence of TYLCV infection (Medina-Puche et al., 2020) or when coexpressed with SlCOPIβ (Figure 1D), we reasoned that the expression of COPI components, particularly SlCOPIβ, might be stimulated during TYLCV infection. As expected, we noticed 2.5- and 3.6-fold increases of accumulated *SlCOPIβ* transcripts at 12 days and 24 days post-inoculation (dpi) of TYLCV in TYLCV-inoculated plants compared with mock-inoculated plants, respectively (Supplemental Figure S2). To better assess the biological significance of the C4-SlCOPIβ interaction *in vivo*, we overexpressed *SlCOPIβ* in tomato using a Potato virus X (PVX) vector, which is commonly used to overexpress genes in tomato (Kong et al., 2013). PVX or PVX-SlCOPIβ was launched by agroinfiltration. At 8 days post-agroinfiltration (dpai), a ∼2.5-fold increase of accumulated *SlCOPIβ* transcripts was observed in PVX-SlCOPIβ plants in comparison to that in PVX plants (Figure 2A). No growth defects were found in PVX-SlCOPIβ plants at 16 dpai (Supplemental Figure S3A and S3B). We next infected PVX- and PVX-SlCOPIβ-treated plants with TYLCV at 8 days after PVX infiltration, and checked symptom development at 24 days after TYLCV infection. Although severe symptoms were present in all plants, PVX-SlCOPIβ plants showed stronger symptoms than that of PVX plants (Figure 2B). In addition, 2.5-fold more viral DNA was accumulated in PVX-SlCOPIβ plants than that in PVX plants (Figure 2C).

**Figure 2.**
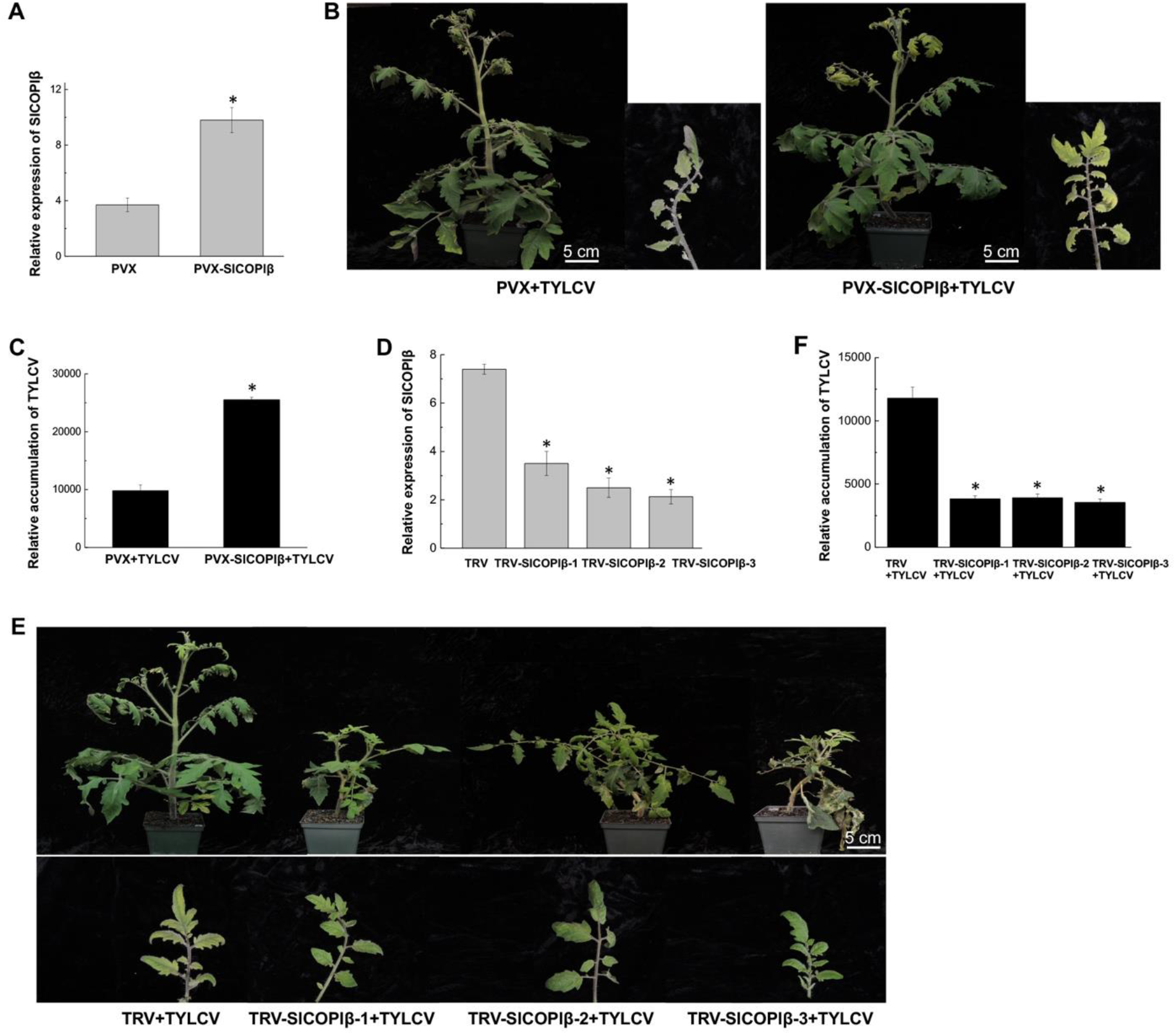
SlCOPIβ plays a critical role in TYLCV infection. (A) Quantitative RT-PCR results showing the relative levels of *SlCOPIβ* transcripts in tomato plants at 8 days after the treatment with PVX or PVX-SlCOPIβ. *SlActin* was used as an internal reference. Values represent the mean ± SD (standard deviation) (n=3 biological replicates). Asterisk indicates a statistically significant difference (*: p<0.05) based on a one-way ANOVA test. (B) Symptoms on TYLCV-infected PVX and PVX-SlCOPIβ plants. Plants and newly emerged leaves were photographed 24 dpi. Bars: 5cm. (C) Quantitative PCR showing the accumulated viral genomic DNA in TYLCV-infected PVX or PVX-SlCOPIβ tomato plants at 24 dpi. *: p < 0.05 (one-way ANOVA test). (D) Quantitative RT-PCR showing the relative levels of *SlCOPIβ* transcripts in TRV and TRV-SlCOPIβ tomato plants at 10 dpai as (A). (E) TYLCV-caused symptoms in control or *SlCOPIβ*-silenced plants. Leaf images were taken at 24 days after TYLCV infection. Bar: 5cm. (F) The viral genomic DNA in systemic leaves of control or *SlCOPIβ*-silenced plants infected with TYLCV measured by qPCR at 24 dpi. *: p < 0.05 (one-way ANOVA test).

We further tested the effect of *SlCOPIβ* knockdown on TYLCV infection in tomato plants by using a Tobacco rattle virus (TRV)-mediated, virus-induced gene silence (VIGS) approach (Liu et al., 2002). To monitor VIGS progress, we also included *phytoene desaturase* (PDS) gene as a reporter (Zhao et al., 2021). TRV, TRV-PDS, and TRV-SlCOPIβ were launched by agroinfiltration. The *SlCOPIβ* expression was downregulated in the TRV-SlCOPIβ lines by 50%-70% at 10 dpai compared to the TRV control (Figure 2D). At 16 dpai, the average height of *SlCOPIβ*-silenced plants was 15 cm, significantly shorter than that of control plants at 23 cm (Supplemental Figure S4A and S4B). Leaf chlorosis and vein necrosis were also found in the *SlCOPIβ-*silenced lines at 30 dpai (Supplemental Figure S4C), agreeing with the critical role of *COPIβ* in Arabidopsis growth (Sánchez-Simarro et al., 2020).

TRV and TRV-SlCOPIβ plants were next inoculated with TYLCV at 10 days after launching TRV, when there were no obvious differences on plant height between them. TYLCV infection caused weaker symptoms in TRV-SlCOPIβ plants compared to control plants at 24 dpi (Figure 2E). Moreover, accumulated viral DNA decreased ∼70% in *SlCOPIβ*-silenced plants than in TRV plants (Figure 2F), suggesting that *SlCOPIβ* serves as a proviral factor for TYLCV infection.

### Coexpression of SlCOPIβ and C4 proteins lead to the C4 localization at chloroplasts and the perinuclear clustering of chloroplasts

Although C4-YFP fluorescence signals were observed primarily in the PM, PD, and weakly in chloroplasts when expressed alone (Figure 1A; Medina-Puche et al., 2020), we noticed that the reconstituted YFP signals were enriched at chloroplasts, which clustered around the nucleus when SlCOPIβ-YFP^N^ and C4-YFP^C^ were coexpressed (Figure 1D). It has been reported that chloroplasts assemble around the nucleus and C4 relocates to chloroplasts during infections of various geminiviruses or the expression of genimivirus protein Rep (Ding et al., 2019). We confirmed that, indeed, chloroplasts clustered around the nucleus and a pool of C4 was detected in the nucleus-surrounding chloroplasts during TYLCV infection (Figure 3A and Supplemental Figure S5). To further clarify the effect of SlCOPIβ on the relocalization of C4 and perinuclear clustering of chloroplasts, we coexpressed FLAG-tagged SlCOPIβ (FLAG-SlCOPIβ) and C4-YFP in *N. benthamiana* cells. We confirmed a strong YFP signal at chloroplasts, which surrounded the nucleus in ∼63% of cells (n=50), in comparison to that of ∼10% of cells with the C4 protein alone (Figure 3B and 3C). Note that we counted cells if more than 4 chloroplasts surrounded the nucleus.

**Figure 3.**
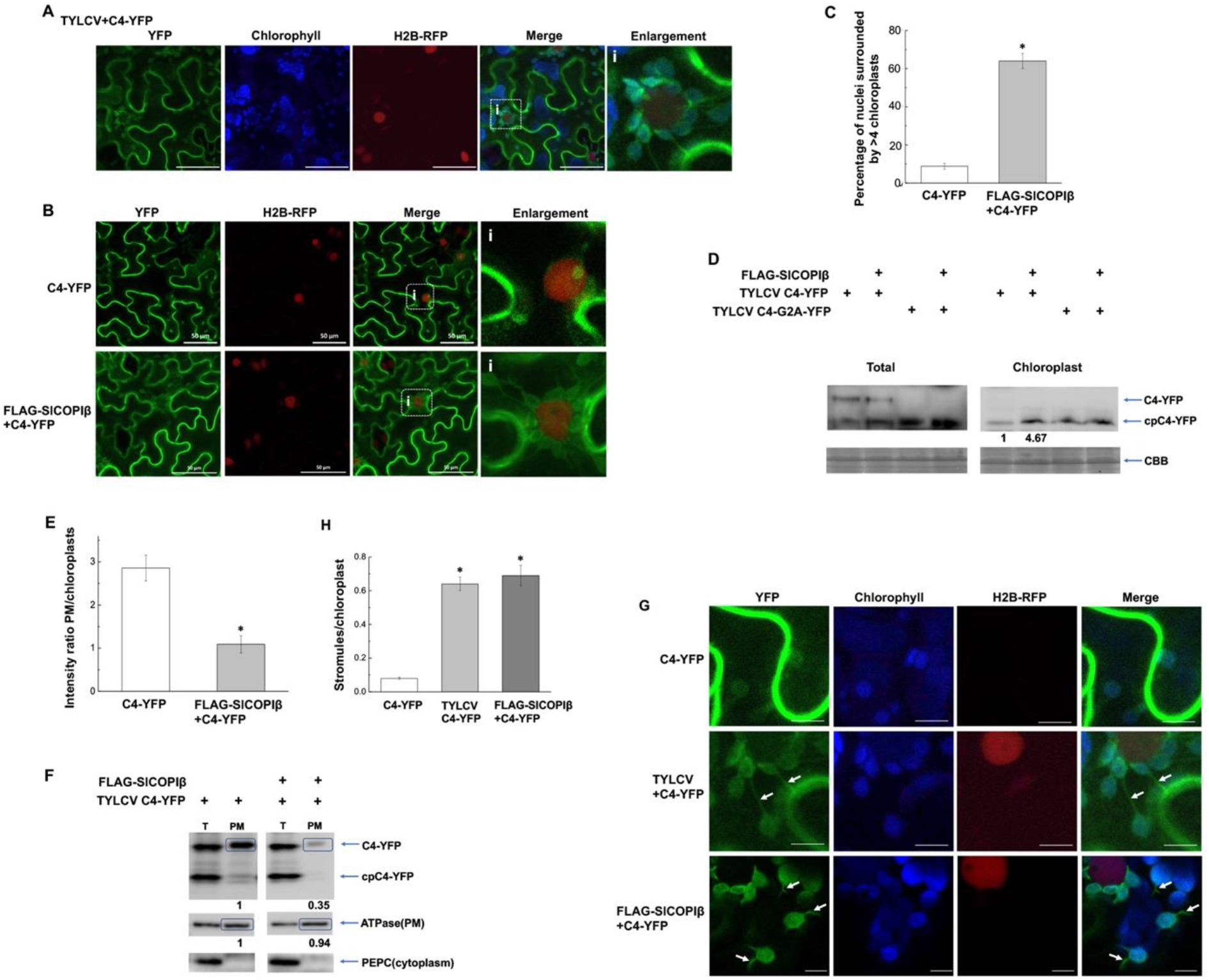
The effect of SlCOPIβ on the enrichment of C4 protein at chloroplasts. (A) Localization of C4-YFP in TYLCV-infected H2B transgenic *N. benthamiana* cells. The rectangle indicates the enlarged region on the right. Bars: 50 μm. (B) Localization of C4-YFP in the absence or presence of the FLAG-SlCOPIβ in H2B transgenic *N. benthamiana* cells. Bars: 50 μm. (C) Percentage of cells with nuclei that are surrounded by four or more chloroplasts in cells expressing C4-YFP with or without SlCOPIβ. Data are means ± SD (n=3). Asterisk indicates a statistically significant difference (*: p<0.05) based on a one-way ANOVA test. (D) Western blot showing total lysate- and the chloroplast fraction-localized C4 protein that was transiently expressed in *N. benthamiana* with or without SlCOPIβ. Total: total lysates. Chloroplast: the isolated chloroplast fraction. CBB: rubisco large subunit as stained by Coomassie brilliant blue. Numbers below blots indicate relative intensity. C4-YFP: the PM-localized full-length C4-YFP; cpC4-YFP: the chloroplast-localized, processed form of C4-YFP. (E) YFP intensity ratios of the PM- and chloroplast-localized C4-YFP without or with SlCOPIβ (from images shown in B). The YFP signal intensity was quantified by using ImageJ. Bars represent SD of n=6. *: p < 0.05 (one-way ANOVA test). (F) Membrane fractionation assays showing PM-accumulated C4-YFP with or without SlCOPIβ. C4-YFP: the PM-localized full-length C4-YFP; cpC4-YFP: the chloroplast-localized, processed form of C4-YFP. PEPC and ATPase showing cytoplasm or PM fraction, respectively. (G) Stromules (pointed by white arrows) were observed in H2B transgenic *N. benthamiana* cells coexpressing C4-YFP and FLAG-SlCOPIβ or during TYLCV infection. Bars: 10 μm. (H) Quantification of the ratio of stromules/chloroplasts from experiments described in (G).

We next isolated chloroplasts and tested the processed form of C4-YFP, cpC4-YFP, in chloroplasts where the cTP signal was cleaved. Full-length C4-YFP was only detected in the total fraction but not the chloroplast fraction. As shown in the total fraction, additional bands, which were possible degradation products from the PM pool of C4-YFP (Figure 3D), as previously reported in Medina-Puche et al. (2020). However, cpC4-YFP, which was ∼33 kDa, accumulated in the chloroplasts fraction. Interestingly, a 4-fold increase of accumulated cpC4-YFP was achieved when C4-YFP was coexpressed with SlCOPIβ compared to the expression of C4-YFP alone (Figure 3D). In contrary, the amount of the non-myristoylable mutant C4-G2A-YFP, which exclusively localized in chloroplasts (Figure 1E), was not affected by the coexpressed SlCOPIβ (Figure 3D, the chloroplast fraction).

We made two measurements to demonstrate that the increase of chloroplast-localized C4-YFP was at the expense of the PM-localized C4-YFP. We measured the intensity of YFP signal at the PM and chloroplasts, and calculated the ratio of the PM- and chloroplast-accumulated C4-YFP, following the procedure as reported in Medina-Puche’s (2020). As shown in Figure 3E, the PM/chloroplast C4-YFP ratio significantly decreased by ∼60% in the presence of SlCOPIβ compared to that without SlCOPIβ. We also performed a cell fractionation assay to enrich the PM fraction (Jacob et al., 2021). As shown in Figure 3F, the PM-localized full-length C4-YFP decreased by 64% when co-expressed with SlCOPIβ, in comparison to that without SlCOPIβ, confirming that the chloroplast-localized C4-YFP was redistributed from the PM.

In Figures 1D and 3B, we tagged C4 with full-length or C-terminus YFP. To exclude a possible contribution from YFP, we tagged both C4 and SlCOPIβ with FLAG tag. When FLAG-C4 and FLAG-SlCOPIβ were coexpressed, chloroplasts also clustered around the nucleus in ∼68% of cells (n=50 cells) (Supplemental Figure S6A and S6B). However, the alignment of chloroplasts along nuclei was found in <10% of cells when FLAG-C4 or FLAG-SlCOPIβ was expressed alone. In addition, coexpressing C4-G2A and SlCOPIβ did not promote chloroplasts perinuclear clustering. To rule out the possibility that any geminiviral protein may induce perinuclear clustering of chloroplasts via the host’s COPI pathway, we included TYLCV V2 protein, a viral silencing suppressor, as a control. Coexpression of FLAG-V2 and FLAG-SlCOPIβ had no significant effect on the localization of chloroplasts (n=50, Supplemental Figure S6A and S6B). These results suggested that the perinuclear clustering of chloroplasts is specifically promoted by the C4-SlCOPIβ interaction and is related to the increased expression of *SlCOPIβ*.

It is increasingly clear that chloroplasts establish physical contact with the nucleus, by clustering around the nucleus and producing extensive tubular stroma-filled protrusions called stromules (Kumar et al., 2018; Kwok and Hanson, 2004; Caplan et al., 2015). Here, we also found stromules during TYLCV infection or when C4-YFP was coexpressed with SlCOPIβ (Figure 3G). Following early reports (Kumar et al., 2018; Caplan et al., 2015), we quantified stromules from 30 epidermal cells and calculated the stromule/chloroplast ratio. As shown in Figure 3H, the ratio increased from 0.1 in control cells to 0.6 during TYLCV infection or 0.65 when C4 and SlCOPIβ were coexpressed. It needs to note that C4-YFP was also detected in these stromules.

Because TYLCV C4 is reported to block SA biosynthesis in chloroplasts (Medina-Puche et al., 2020), we measured the expression of two SA-related genes in *N. benthamiana* plants expressing C4 alone or with SlCOPIβ: *Isochorismate Synthase 1* (*ICS1*) that encodes a key enzyme in SA biosynthesis and *Pathogenesis Related protein 1* (*PR-1a*) that is regulated by SA. As shown in Supplemental Figure S7, the coexpression of C4 and SlCOPIβ resulted in lower expression levels of both *ICS1* and *PR-1a* genes compared to mock plants, in sharp contrast to those expressing C4 alone. These results are consistent with the published results (Medina-Puche et al., 2020) and with our data that TYLCV infection was promoted when *SlCOPIβ* was overexpressed but inhibited when the gene expression of *SlCOPIβ* was knocked down (Figure 2).

### COPI components contribute to the perinuclear clustering of chloroplasts and TYLCV infection

COPI vesicles are composed of seven subunits: α, β, β’, γ, δ, ε, and ζ (Waters et al., 1991; Arakel and Schwappach, 2018; Bethune and Wieland, 2018). To test whether other COPI components are also involved in TYLCV infection, we selected two COPI members: SlCOPIδ is a component of COPI adaptor-like subcomplex, the same subcomplex with SlCOPIβ; SlCOPIε is from COPI coat-like subcomplex (Ritzenthaler et al., 2002). We first knocked down the gene expression of *SlCOPIδ* or *SlCOPIε* in tomato plants. *SlCOPIδ*- or *SlCOPIε*-silenced plants were shorter than control plants, similar to that of TRV-SlCOPIβ plants (Supplemental Figure S8A and S8B). Similar growth phenotype, such as leaf chlorosis and vein necrosis, were also found in *SlCOPIδ*- or *SlCOPIε*-silenced plants (Supplemental Figure S8C), indicating that the COPI components play an important role in plant growth.

*SlCOPIδ*- or *SlCOPIε*-silenced plants were then inoculated with TYLCV, and infection progress was monitored over time. Similar to *SlCOPIβ*-silenced plants, *SlCOPIδ*- or *SlCOPIε*-silenced plants infected with TYLCV developed weaker symptoms than that in control plants (Figure 4A and 4B). Viral DNA accumulated at much lower levels in the silenced plants than in control plants, only 20-25% of that in TRV-treated plants (Figure 4C). At 20 days after TYLCV inoculation, only ∼2 out of the 15 *SlCOPIδ*- or *SlCOPIε*-silenced plants showed minor symptoms, but ∼13 control plants showed strong symptoms (Figure 4D). Although ∼45% of the silenced lines developed symptoms by 28 days, the silenced plants showed much weaker symptoms, similar to *SlCOPIβ*-silenced plants. These results suggest that the host COPI complex, not just SlCOPIβ, is involved in TYLCV systemic infection.

**Figure 4.**
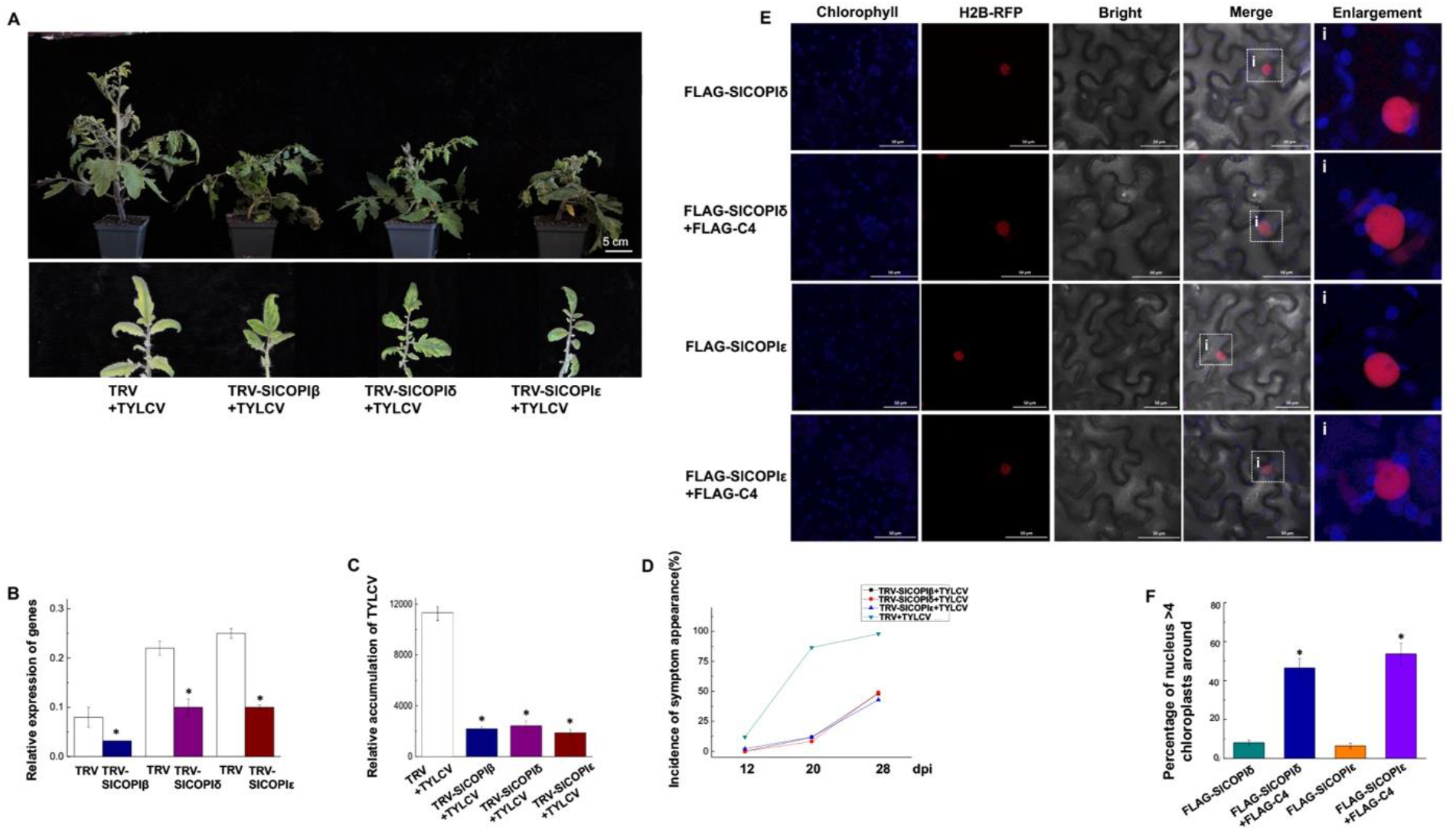
Tomato COPI pathway is involved in TYLCV infection. (A) Symptoms in TYLCV-infected TRV-, TRV-SlCOPIβ-, -SlCOPIδ- or -SlCOPIε-treated plants at 24 dpi. Bar: 5cm. (B) The relative levels of accumulated *SlCOPIβ*, *SlCOPIδ*, or *SlCOPIε* transcripts in control or knockdown plants were measured by qRT-PCR at 10 dpai as Figure 2A. (C) The accumulated TYLCV genomic DNA in upper leaves as measured by qPCR at 24 dpi. *: p < 0.05 (one-way ANOVA test). (D) Disease incidence in control or knockdown plants. The percentages of TYLCV infected plants were determined with symptoms and qPCR at the indicated time points. 15 tomato plants were inoculated with TYLCV in each treatment. (E) Chloroplast localizations upon transient expression of FLAG-SlCOPIδ or -SlCOPIε in the absence or presence of FLAG-C4. Bars: 50 μm. (F) Percentage of cells with the nucleus surrounded by four or more chloroplasts expressing FLAG-SlCOPIδ or -SlCOPIε alone or with FLAG-C4. Asterisk indicates a statistically significant difference (*: p<0.05) based on a one-way ANOVA test.

To test the effect of COPI components on chloroplasts, we then investigated the localization of chloroplasts when C4 was coexpressed with SlCOPIδ or SlCOPIε. In 48-57% of cells coexpressing FLAG-C4 and FLAG-SlCOPIδ or -SlCOPIε, chloroplast fluorescence signal was observed around the nuclei (Figure 4E and 4F), suggesting that COPI components play important role for the clustering of chloroplasts around the nucleus and, in turn, affect viral infection.

The fact that the interaction sites of SlCOPIβ and C4 were primarily at chloroplast and weakly at the PM (Figure 1D) promoted us to check the localization of SlCOPIβ in the presence of C4 or during TYLCV infection. We first transiently expressed SlCOPIβ-YFP in H2B transgenic *N. benthamiana* plants by agroinfiltration and noticed that a very strong YFP signal was found primarily at chloroplasts in TYLCV-infected cells (Figure 5A). The SlCOPIβ-YFP was also enriched in chloroplasts when coexpressed with FLAG-C4 (Figure 5A), even though weaker than that in TYLCV-infected cells. Accompanied by the increased chloroplast localization of SlCOPIβ-YFP, ∼50% of cells (n=30) had clustered chloroplasts around the nucleus (Figure 5A and 5B). These results are consistent with our BiFC results that the reconstituted YFP signal due to the C4-SlCOPIβ interaction was enriched in chloroplasts (Figure 1D).

**Figure 5.**
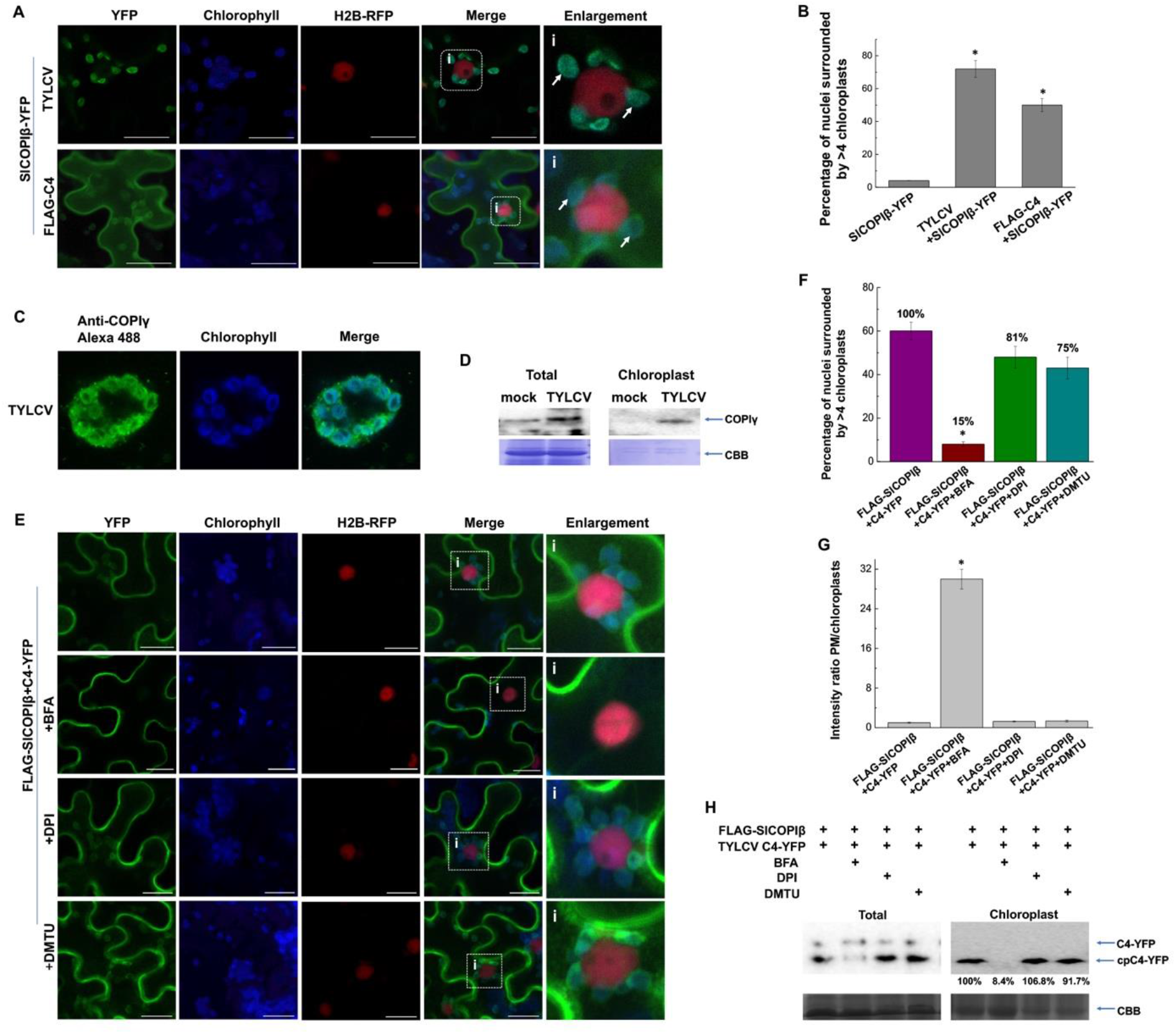
The perinuclear clustering of chloroplasts induced by coexpression of SlCOPIβ and C4 requires the COPI pathway. (A) Localization of SlCOPIβ-YFP during TYLCV infection or when coexpressed with FLAG-C4 in H2B transgenic *N. benthamiana* cells. White arrows point to chloroplasts. The rectangle indicates the enlarged region on the right. Bars: 20 μm. (B) Percentage of cells with nuclei that are surrounded by four or more chloroplasts in leaves expressing SlCOPIβ-YFP as (A). *: p<0.05, (one-way ANOVA test). (C) Immunofluorescence microscopic images showing the localization of tomato COPIγ in TYLCV-infected tomato protoplasts. Localization of COPIγ was determined by using a polyclonal antibody recognizing COPIγ and a secondary anti-rabbit antibody conjugated to Alexa Fluor 488. (D) Western blot showing SlCOPIγ in total and chloroplast fraction prepared from mock- or TYLCV-infected tomato plants. Total lysates or the isolated chloroplast fraction were incubated with anti-COPIγ antibody-bound Sepharose beads at 4°C overnight and SlCOPIγ was detected by using the polyclonal anti-COPIγ antibody. Total: total lysates. Chloroplast: the isolated chloroplast fraction. CBB: Rubisco large subunit as stained by Coomassie brilliant blue. (E) Localization of C4-YFP following treatment with BFA (20 μM, 6 h treatment), DPI (50 μM, 2 h), or DMTU (5 mM, 2h) in the presence of FLAG-SlCOPIβ. The rectangle indicates the enlarged region. Bars: 20 μm. (F) Percentage of cells with nuclei that are surrounded by four or more chloroplasts in cells coexpressing C4 and SlCOPIβ and further treated with BFA, DPI or DMTU. The percentage of coexpressed C4 and SlCOPIβ was set as 100%. *: p<0.05, (one-way ANOVA test). (G) Intensity ratios of the PM/chloroplast-localized C4-YFP when coexpressed with SlCOPIβ upon different chemical treatment (from E) as Figure 3E. (H) Western blot showing total- and chloroplast-localized C4 in the presence of SlCOPIβ with BFA, DPI, or DMTU treatment. Cell fraction and Western were performed as Figure 3D.

We next prepared protoplasts from TYLCV-infected tomato plants, and tested localization of a COPI component, COPIγ, using a specific antibody in an immunofluorescence assay (Hurný et al., 2020; Singh et al., 2018; Nagel et al., 2017). In TYLCV-infected tomato cells, strong signals were concentrated at patches that colocalized with chloroplasts (Figure 5C, Supplemental Figure S9). We additionally detected COPIγ in the chloroplast fraction by using western blotting. The COPIγ band was only detected in the chloroplast fraction of TYLCV-infected tomato protoplasts but not control plants (Figure 5D). We also observed an increased accumulation of COPIγ in total lysate, similar to the increased accumulation of SlCOPIβ transcript during TYLCV infection (Supplemental Figure S2). Collectively, our results suggested that COPI vesicles localized to chloroplasts, not only SlCOPIβ, in the presence of C4 or during TYLCV infection.

To further verify the role of the COPI pathway in the relocalization of C4 and the clustering of chloroplasts in the perinuclear region, we tested the subcellular localization of C4 when coexpressed with SlCOPIβ upon treatment with Brefeldin A (BFA). BFA blocks the activity of GBF1, and in turn, inhibits the formation of COPI vesicles (Ritzenthaler et al., 2002). At 6 hours post-treatment with 20 μg/mL BFA, two changes were readily noticed: the percentage of cells with nucleus-surrounding chloroplasts decreased to <10% (n=30 cells) (Figure 5E and 5F) and the C4-YFP signals were no longer dominantly observed in chloroplasts, but were, rather, dominant in the PM. The intensity ratio of PM- and chloroplast-localized YFP signal increased ∼30-fold with BFA treatment (Figure 5G), and also cpC4-YFP was barely detected in the isolated chloroplast fraction in the presence of BFA (Figure 5H), suggesting that COPI complexes are involved in transporting C4 to chloroplasts and the enrichment of chloroplasts around the nucleus.

Perinuclear clustering of chloroplasts can be induced by treatment with flg22, an elicitor of pattern-triggered immunity (PTI), or by an exogenous application of ROS, such as H_2_O_2_ (Ding et al., 2019; Gómez-Gómez et al., 1999). It was further shown that ROS was necessary in the flg22-induced perinuclear clustering of chloroplasts, because treatment of the NADPH-oxidase inhibitor diphenyleneiodonium (DPI) or the ROS scavenger dimethylthiourea (DMTU) dramatically decreased perinucleus-localized chloroplasts (Ding et al., 2019). We tested whether accumulation of ROS is also required for SlCOPIβ/C4-mediated perinuclear clustering of chloroplasts. In *N. benthamiana* leaves that have been agroinfiltrated with both SlCOPIβ and C4, we further infiltrated DPI or DMTU solution and harvested tissues 2 hours later. Even though the treatment by DPI or DMTU blocked the flg22-induced perinuclear clustering of chloroplasts (Supplemental Figure S10A and S10B), neither chemical had observable effects on the number of chloroplasts surrounding nuclei in the presence of FLAG-SlCOPIβ and C4-YFP (Figure 5E) or FLAG-SlCOPIβ and FLAG-C4 (Supplemental Figure S11A and S11B). Conversely, treatment with BFA substantially affected the number of chloroplasts that were aligned around nuclei (Figure 5E and 5F), suggesting that ROS are not required for chloroplasts to surround the nucleus as induced by coexpression of SlCOPIβ and C4. To our surprise, the BFA treatment also mitigated the flg22-mediated perinuclear enrichment of chloroplasts, even though not as effective as DPI or DMTU in the concentrations we used (Supplemental Figure S10A and S10B).

### The COPI pathway is broadly involved in chloroplast-targeting of C4/AC4 proteins and infections of geminiviruses

It has been previously reported that besides TYLCV C4, both of C4 from the curtovirus BCTV and AC4 from the bipartite begomovirus EACMV have two targeting signals: an N-myristolation motif and a cTP (Medina-Puche et al., 2020). It was further shown that all three viral proteins can be enriched at chloroplasts upon treatment with flg22 (Medina-Puche et al., 2020). Given that the three viral proteins share similar features, we hypothesized that the COPI pathway is involved in the chloroplast targeting of BCTV C4 and EACMV AC4, as well as infections of both viruses, similar to those of TYLCV.

As expected, a very strong fluorescence signal of BCTV C4- or EACMV AC4-GFP was found in the PM and a weak signal in the chloroplasts in almost all cells (n=30) (Figure 6A), and the non-myristoylable G2A mutant accumulated in chloroplasts exclusively (n=30) (Figure 6A), similar to TYLCV C4 and as previously reported (Medina-Puche et al., 2020). In a co-IP assay as shown in Figure 6B, FLAG-SlCOPIβ was pulled down with TYLCV C4, BCTV C4, or EACMV AC4, but not TYLCV C4-G2A. Similar results were achieved in the Y2H analysis where cells expressing SlCOPIβ and TYLCV C4, BCTV C4 or EACMV AC4 grew well on the selection medium, but not cells coexpressing G2A mutant and SlCOPIβ (Figure 6C).

**Figure 6.**
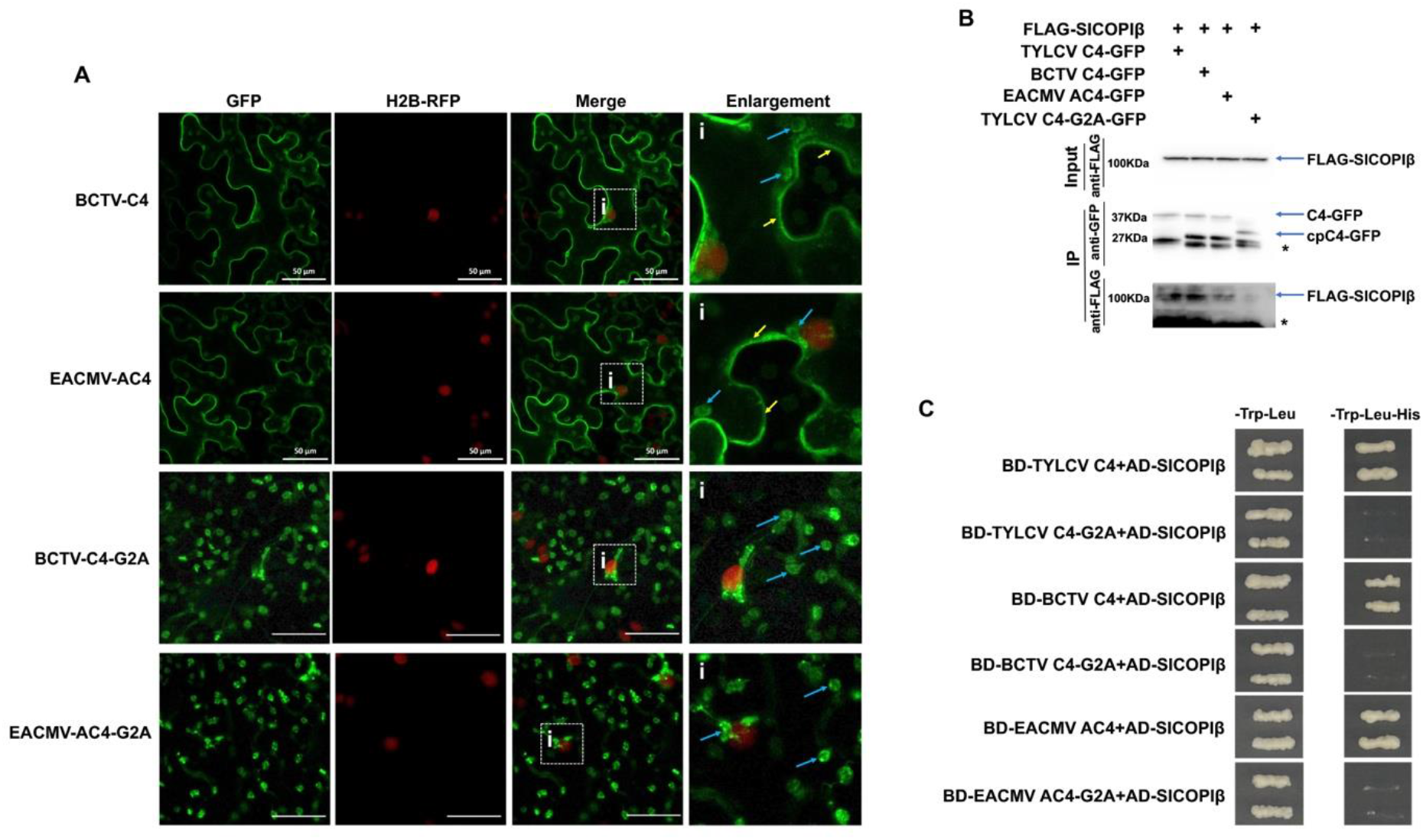
C4/AC4 proteins from other geminiviruses interact with SlCOPIβ. (A) Localization of BCTV C4 or EACMV AC4 in H2B transgenic *N. benthamiana* cells. Yellow arrows point to the PM; blue arrows point to chloroplasts. The rectangle indicates the enlarged region on the right. Bars: 50 μm. (B) The interaction between SlCOPIβ and TYLCV C4, BCTV C4, or EACMV AC4 was tested by a co-IP assay. (C) Interactions between SlCOPIβ and WT C4/AC4 or G2A mutants from different geminiviruses were tested using Y2H as Figure 1F.

We next tested whether C4/AC4 from BCTV/EACMV could be localized to chloroplasts when coexpressed with SlCOPIβ. As expected, a very strong fluorescence signal of BCTV C4- or EACMV AC4-GFP was found in the PM and a weak signal in the chloroplasts in almost all cells (n=30) when expresses alone (Figure 7A), similar to TYLCV C4 and as previously reported (Medina-Puche et al., 2020). However, when coexpressed with FLAG-SlCOPIβ, both were primarily detected at chloroplasts, based on confocal microscopy (Figure 7A) or cell fraction assay (Figure 7B). Similar to TYLCV C4-YFP (Figure 3E), the ratio of PM- and chloroplast-localized BCTV C4-GFP or EACMV AC4-GFP significantly decreased in the presence of SlCOPIβ compared to that in the absence of SlCOPIβ (Figure 7C), suggesting that the accumulation of C4/AC4-GFP at chloroplasts increased when coexpressed with SlCOPIβ. In contrary, the G2A substitution in BCTV C4 and EACMV AC4 did not have any effect on chloroplast distribution when combined with SlCOPIβ (Figure 7D and 7E). In addition, we found that chloroplasts were enriched around the nucleus in ∼65% cells (n=30) coexpressing FLAG-SlCOPIβ and BCTV C4-GFP or EACMV AC4-GFP (Figure 7E), similar to that of coexpression of FLAG-SlCOPIβ and TYLCV FLAG-C4 (Supplemental Figure S6B).

**Figure 7.**
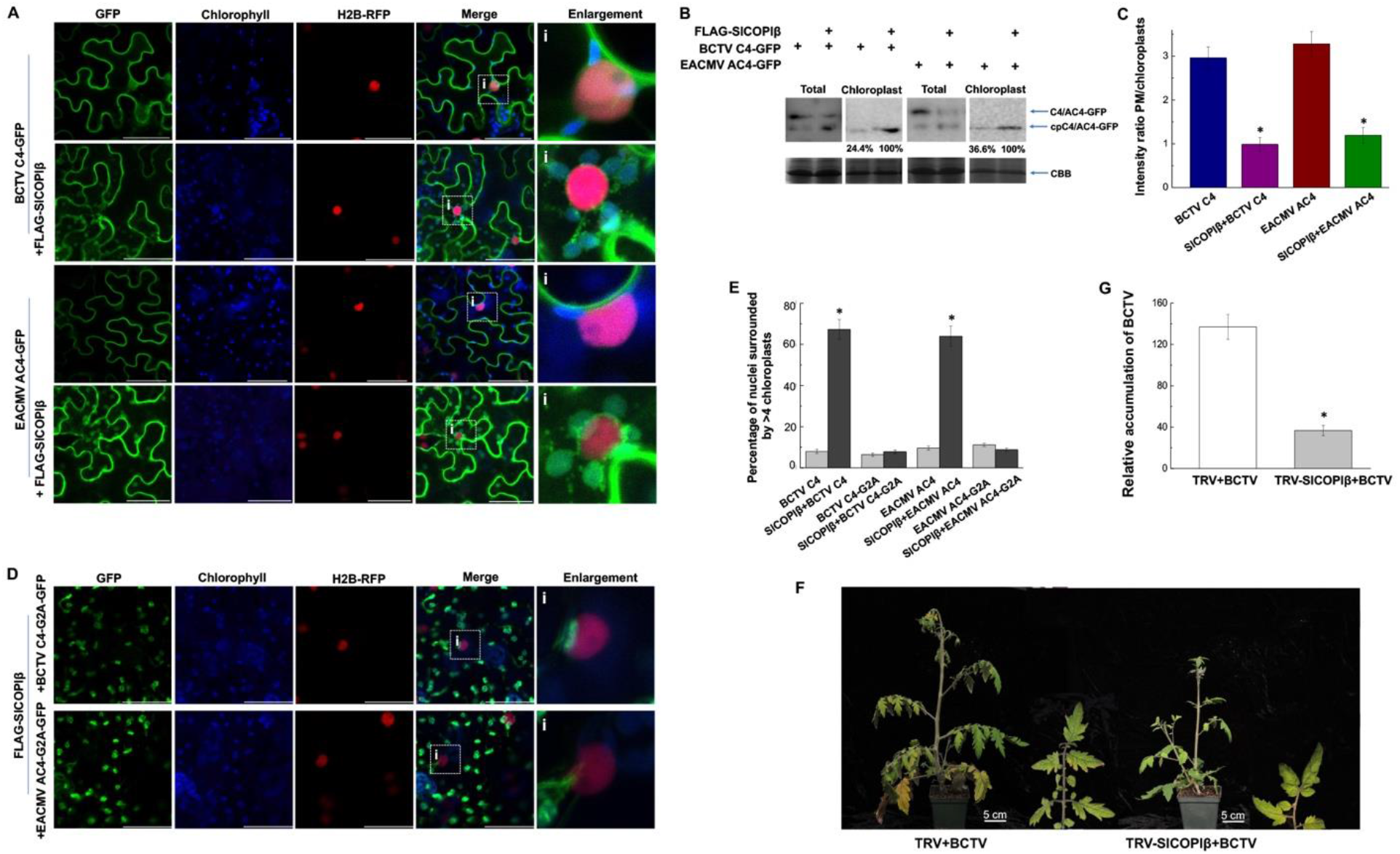
SlCOPIβ is involved in the redistribution of C4/AC4 proteins of other geminiviruses to chloroplasts. (A) Localization of BCTV C4-GFP or EACMV AC4-GFP in the absence or presence of SlCOPIβ. Bars: 50 μm. (B) Western blot showing total lysate- and the chloroplast-localized BCTV C4/EACMV AC4 in the absence or presence of SlCOPIβ. Cell fractionation and western blot were done as in Figure 3D. (C) Intensity ratios of PM- and chloroplast-localized BCTV C4-GFP and EACMV AC4-GFP in the absence or presence of SlCOPIβ. The intensity of GFP signals was quantified by using ImageJ. *: p < 0.05 (one-way ANOVA test). (D) Localization of G2A mutants of BCTV C4 or EACMV AC4 in the presence of SlCOPIβ in *N. benthamiana* cells. Bars: 50 μm. (E) Percentage of cells with nuclei that are surrounded by four or more chloroplasts in cells coexpressing SlCOPIβ and WT or G2A mutants of geminiviruses C4/AC4 proteins. Data the mean ± SD (n=3), *: p < 0.05 (one-way ANOVA test). (F) Symptoms caused by BCTV infection in TRV- or TRV-SlCOPIβ-treated plants at 30 dpi. Bar: 5cm. (G) The accumulated BCTV viral DNA in systemic leaves as measured by qPCR at 30 dpi. *: p < 0.05 (one-way ANOVA test).

To further test the role of BCTV C4-SlCOPIβ interaction in BCTV infection, *SlCOPIβ*-silenced plants were inoculated with BCTV and infection progress was monitored. BCTV-infected *SlCOPIβ*-silenced tomato plants developed mild symptoms (Figure 7F) with lower levels of accumulated viral DNAs than that of control plants (Figure 7G), suggesting a broad proviral role of SlCOPIβ in geminivirus infections.

## Discussion

Although the primary function is photosynthesis, chloroplasts are also the site for producing defense hormones, such as JA precursors, SA, and ROS (Chan et al., 2016). In response to various stresses, chloroplasts cluster around the nucleus, and may deliver defense signaling molecules, such as SA and H_2_O_2_, to the nucleus via stromules (Kumar et al., 2018; Kwok and Hanson, 2004). These defense signals induce the expressions of a cluster of host genes and/or the programmed cell death (Kumar et al., 2018; Kwok and Hanson, 2004). To combat host innate immunity, geminiviruses-encoded C4/AC4 proteins shuttle from the PM to chloroplasts where they interfere with SA production, sabotaging SA-regulated defense responses and promoting viral infection (Medina-Puche et al., 2020). However, mechanism(s) by which C4/AC4 proteins transport to chloroplasts are unknown. We have identified an unexpected role of the COPI pathway in transporting C4/AC4 proteins to chloroplasts. We demonstrated that TYLCV C4-YFP localizes primarily in the PM/PD when expressed alone (Figure 1A and Supplemental Figure S1A), but was highly enriched in the perinuclear localized chloroplasts in the presence of SlCOPIβ in *N. benthamiana* cells (Figure 3B). We further showed that a COPI inhibitor, BFA, disrupted chloroplast localization of C4 and perinuclear clustering of chloroplasts as promoted by the coexpression of C4 and SlCOPIβ (Figure 5E and 5F). Consistent with the role of COPI pathway in C4/AC4 targeting, knocking down gene expression of SlCOPIβ, δ, or ε inhibited TYLCV and BCTV infection, but overexpression of SlCOPIβ promoted TYLCV infection (Figures 2, 4A, 4C, 4D, 7F and 7G). Our results, therefore, revealed that COPI pathway played a novel role in the trafficking of geminiviral proteins to chloroplasts during geminivirus infections.

Our data also demonstrated that the interaction between C4/AC4 and SlCOPIβ is required to induce the enrichment of chloroplasts at the perinuclear region (Figures. 1C, 1D, 6C). The non-myristoylable mutants of C4 proteins from TYLCV and BCTV or AC4 from EACMV, C4/AC4-G2A, localizes to chloroplasts exclusively without distribution at the PM/PD (Figure 1E and 7D; Rosas-Diaz et al., 2018). Here, we found that the G2A mutation failed to interact with SlCOPIβ (Figures 1F, 1G and 6C), and overexpression of both C4/AC4-G2A and SlCOPIβ triggered perinuclear clustering of chloroplasts only in ∼10% of cells, a sharp decrease from the ∼65% of cells that coexpressing WT C4/AC4 and SlCOPIβ (Supplemental Figure S6B and Figure 7E). However, we cannot totally rule out the possibility that the G2A mutants interact with SlCOPIβ in plant because the G2A mutant did not localize to the PM and did not have a chance to encounter SlCOPIβ.

While it is clear that trafficking of C4/AC4 to the PM requires host N-myristoyl transferase, it is unclear how C4/AC4 proteins are transported to chloroplasts (Medina-Puche et al., 2020). Our data showed that TYLCV C4-YFP was detected primarily in chloroplasts and weakly in PM when coexpressed with FLAG-SlCOPIβ (Figure 3B), however, C4-YFP was only detected in the PM but not in chloroplasts when treated with BFA (Figure 5E), suggesting that SlCOPIβ and the COPI pathway are involved in transporting C4 protein to chloroplasts. We provided two lines of evidence to further demonstrate that the increased pool of processed C4 at chloroplasts might be redistributed from the PM. In a cell fractionation assay, significantly increased C4 was detected at chloroplasts and the PM-localized full-length C4-YFP decreased 64% (Figure 3D and 3F). We also measured the signal intensity of C4-YFP and calculated the ratio of PM/chloroplasts signal intensity and found a significant decrease of the ratio in the presence of both C4-YFP and FLAG-SlCOPIβ (Figure 3E).

Our data also suggested that high levels of COPI components are likely needed for targeting C4 to chloroplasts (Figures 3B and 4E). Consistent with this notion, a 2.5-fold to 3.6-fold increase of SlCOPIβ transcripts (Supplemental Figure S2) or an increased protein levels of the endogenous SlCOPIγ (Figure 5D) during TYLCV infection were revealed by western blotting. Given several COPI components were similarly involved in the C4 targeting to chloroplasts (Figure 4E), our results suggest the COPI pathway is involved in the C4’s targeting to chloroplasts during viral infection. The COPI pathway is well known for playing a critical role in retrograde trafficking of proteins within the Golgi and from Golgi to the ER (Pimpl et al., 2000). In the past years, several additional functions have been assigned to the COPI pathway (Soni et al., 2009; Wilfling et al., 2014). However, we found that COPI components are also relocalized to chloroplasts in the presence of C4 or during TYLCV infection (Figure 5A and 5C), and play a critical role in the C4 redistribution to chloroplasts (Figure 3), and thus, geminivirus infections (Figure 4A and 7F).

In light of an unconventional role of the COPI pathway, it was previously shown that the COPI pathway is involved in targeting proteins, such as Glycerol-3-phosphate acyltransferase (GPAT4) and adipose triglycerise lipase (ATGL), from the ER to lipid droplet (LD) (Soni et al., 2009; Wilfling et al., 2014). It is surprising, however, the COPI-mediated protein transport from the ER to LDs is not based on vesicles but membrane bridges connecting LDs and ER membranes (Soni et al., 2009). It has also been reported that COPI and GBF1 are necessary for the transport of Dengue virus (DENV) capsid protein from the ER to LDs where DENV viral particles are assembled (Iglesias et al., 2015). Nevertheless, there has never been a report on a possible role of COPI components in transporting proteins to chloroplasts or in geminivirus infection; therefore, we have identified a novel function of COPI components. Several key questions remain to be answered: Are C4/AC4 proteins cargos of COPI vesicles? How do COPI components/vesicles deliver C4/AC4 to chloroplasts during viral infection? It is also unclear at which organelle COPI components/vesicles interact and/or pack C4/AC4 proteins, be it the PM, ER, or Golgi. It has been proposed that C4/AC4 is first transported to the PM after and then chloroplasts (Medina-Puche et al., 2020). It is possible that COPI vesicles assemble and pack C4/AC4 at the PM. However, the only evidence supporting this possibility is that the reconstituted YFP signal from coexpressed SlCOPIβ-YFP^N^ and C4-YFP^C^ was found at the PM (Figure 1D), we lack direct evidence that COPI components are associated with PM. It is also possible that C4/AC4 proteins interact with COPI components/vesicles at the ER or Golgi and are redirected to chloroplasts without going to the PM. This hypothesis is consistent with our data that the enhanced C4 signal at chloroplasts were accompanied with a decreased C4 accumulation at the PM.

Perinuclear clustering of chloroplasts was previously observed during plant immune responses triggered by flg22, a PTI signaling molecules, or the geminivirus Rep protein, or TMV p50 protein in combination with the host resistance gene N (Kumar et al., 2018; Caplan et al., 2015; Ding et al., 2019; Wilfling et al., 2014). It was further shown that the production of SA or ROS, such as H_2_O_2_, is necessary and sufficient because exogenous application of H_2_O_2_ or SA induced the clustering of chloroplasts surrounding the nucleus and/or the formation stromules (Caplan et al., 2015; Ding et al., 2019). We showed that stromules were formed, perinuclear clustering of chloroplasts occurred, and C4-YFP was detected in stromules during TYLCV infection or TYLCV C4 and SlCOPIβ was coexpressed (Figure 3G). However, the mechanisms by which perinuclear clustering of chloroplasts induced by the flg22 treatment or the coexpression of C4 and SlCOPIβ might be different. Addition of chemical inhibitors of ROS functions blocked the flg22-induced chloroplast clusters (Supplemental Figure S10, Ding et al., 2019), but only slightly affected chloroplast clusters induced by the coexpression of C4 and SlCOPIβ (Figure 5E). Flg22-mediated perinuclear clustering of chloroplast was also significantly inhibited by the BFA treatment (Supplemental Figure S10), however, the inhibitory effect by the BFA treatment might not be direct and requires further investigation. Nevertheless, the mechanisms by which coexpression of C4/AC4 and SlCOPIβ (or other COPI components) induce chloroplasts movement towards the nucleus are unclear. Given that C4/AC4 G2A mutants are primarily localized in chloroplasts but don’t induce perinuclear enrichment of chloroplasts, it is very likely that COPI components contribute to the movement of chloroplasts towards the nucleus by interacting with C4/AC4 proteins.

Importantly, our work demonstrates a critical role of COPI in inhibiting a SA-mediated host defense mechanism during infections of several geminiviruses (Supplemental Figure S7), and knocking down gene expression of several COPI components individually resulted in strong viral resistance (Figures 4A and 7F). Given the fact that geminiviruses contain more than 500 species and cause huge agricultural losses globally, our results open new opportunities in fighting this diverse family of plant pathogens in the future.

## Materials and Methods

### Plant materials and growth conditions

Wild-type (WT) or H2B-RFP (red fluorescent protein was fused at the C terminus of histone 2B) (Martin et al., 2009) transgenic *N. benthamiana* plants was used for agrobacterium infiltration in this study. The highly susceptible cultivar of tomato (*Solanum Lycopersicum*), moneymaker, was used for virus infection. Plants without or with infiltration were grown in a growth chamber at 22-26°C with a 16 h light/8 h dark cycle.

### Plasmid construction

The open reading frame (ORF) of TYLCV C4 was amplified from the TYLCV-infectious clone. The ORFs of SlCOPIβ, SlCOPIδ, and SlCOPIε were cloned from tomato leaf tissue of Moneymaker, using corresponding primers by RT-PCR: 5’-CCATCGATATGGAGAAGTCCTGTTCTTTG-3’ and 5’-GCGTCGACACTGCCTCCCTTCTGCTT-3’ for SlCOPIβ; 5’-ATGGTTGTTCTAGCGGCTTCC-3’ and 5’-CACAACCTGGTAACTCTCCGT-3’ for SlCOPIδ; 5’-ATGGCGTCAACCGGACCAGA-3’ and 5’-AGCAACAGTTTGGACTGCTCT-3’ for SlCOPIε.

FLAG tagged C4, SlCOPIβ, SlCOPIδ, and SlCOPIε ORFs were amplified and inserted into the pCambia1300 binary vector (Zhao et al., 2021) to generate 35S:FLAG-C4, 35S:FLAG-SlCOPIβ, 35S:FLAG-SlCOPIδ, and 35S:FLAG-SlCOPIε for co-IP and co-localization experiments. The full-length SlCOPIβ cDNA was amplified and inserted into the PVX vector to make overexpressing vector. The cDNA fragment of SlCOPIβ (corresponding to amino acids 434 to 534), SlCOPIδ (amino acids 31 to 131) or SlCOPIε (amino acids 76 to 176) were cloned into the TRV vector (Liu et al., 2002) for gene silencing assays.

### Transient expression in *N. benthamiana* plants

YFP- or FLAG-tagged expression plasmids were transformed into *A. tumefaciens* strain GV3101, and the transient gene expression assays were performed as described previously (Zhao et al., 2021). Briefly, the corresponding agrobacterium was grown in LB medium with a combination of antibiotics at 28°C. *Agrobacterium* cultures were collected by centrifuging and suspended in an induction buffer (10 mM MgCl_2_, 100 mM 2-(N-morpholino) ethanesulfonic acid pH 5.7, 150 μM acetosyringone) to an OD_600_=0.5-1 for 2-4 h at room temperature (RT).

For co-localization or gene silencing experiments, *Agrobacterium* cultures haboring two target plasmids were harvested and mixed at 1:1 ratio. The induction mixture was infiltrated into WT or H2B-RFP transgenic *N. benthamiana* plants with a needleless syringe.

### Confocal laser scanning microscopy and co-localization assays

The full-length of TYLCV C4/G2A and SlCOPIβ were amplified using specific primers (Supplemental Table S1) and inserted into the pCambia1300-YFP/-GFP (p1300-YFP/-GFP) vector (Zhao et al., 2021) to generate 35S:C4/G2A-YFP, 35S:C4/G2A-GFP or 35S:SlCOPIβ-YFP. Plasmid expressing YFP- or GFP-tagged proteins were introduced into GV3101 individually and agroinfiltration was performed as above.

The subcellular localization observations were performed using confocal microscopy (ZEISS LSM 880) at 40 hpai. The wavelengths of excitation and emission for YFP and GFP were 488 nm and 497–520 nm, for mRFP were 561 nm and 585–615 nm, and for chloroplast/chlorophyll was 488 nm and 660–720 nm. Signals were sequentially acquired to avoid any possible bleed-through when detecting colocalization.

### Bimolecular fluorescence complementation (BiFC) assay

To make BiFC vectors, the full-length of SlCOPIβ and TYLCV C4 were amplified using specific primers and cloned into the N-terminal fragment of YFP (1-155 aa) or the C-terminal fragment of YFP (156-239 aa), resulting in 35S:SlCOPIβ-YFP^N^ and 35S:C4-YFP^C^. 35S:SlCOPIβ-YFP^N^ and 35S:C4-YFP^C^ were introduced individually into GV3101 and BiFC experiments were performed as described previously (Zhao et al., 2021). 35S:SlCOPIβ-YFP^N^ and 35S:V2-YFP^C^ was used as a negative control. Briefly, the different combinations were co-infiltrated into *N. benthamiana* leaves, and fluorescence signal was observed by using ZEIZZ LSM 880 confocal microscopy at 48 hpai.

### Co-immunoprecipitation assay

For co-IP assay, different combinations of YFP/GFP- and/or FLAG-tagged vectors were expressed in *N. benthamiana* leaves, and performed as previously described (Zhao et al., 2021). Briefly, equal amounts of infiltrated leaf tissues were gently ground with extraction buffer, kept on ice for 20 min, and centrifugated at 21,000xg for 30 min at 4°C. Total lysates were incubated with anti-FLAG beads (Sigma, USA) or monoclonal anti-GFP antibody (Roche, Basel)-bound beads overnight at 4°C. The beads were extensively washed for 5 times. Finally, the equal volume of 2×SDS loading buffer was added. Proteins were detected by a monoclonal anti-FLAG antibody (Sigma, USA) or a polyclonal anti-GFP antibody (GenScript, USA).

### Yeast two-hybrid screen

Library screen was performed as previously described (Zhao et al., 2021) using TYLCV C4 as a bait. The full-length TYLCV C4 was inserted into the pGBKT7 (BD) bait vector and transformed into yeast AH109. The positive interactors were selected from SD/–Leu/–Trp/–His/– Ade selection medium and sequenced.

The Y2H system was used to verify the specific protein-protein interactions and performed as described previously (Zhao et al., 2021). TYLCV C4/C4-G2A, BCTV C4/C4-G2A, EACMV AC4/AC4-G2A and SlCOPIβ ORFs were amplified and inserted into the BD vector or the pGADT7 (AD) vector using the primer pairs: 5’-CGGGATCCATGTCGAAGCGACCAGGCGA-3’ and 5’-CGGGATCCATTTGATATTGAATCATAGAAATAG-3’ for TYLCV C4; 5’-ATGGCGAACCACATCTCCATG-3’ and 5’-CGGGATCCATTTGATATTGAATCATAGAAATAG-3’ for TYLCV C4-G2A; 5’-CACCATGGGCAACCTCATCTCCACG-3’ and 5’-TTAACGCCTTGGCATATGAGTCG-3’ for BCTV C4; 5’-ATGGCCAACCTCATCTCCACG-3’ and 5’-TTAACGCCTTGGCATATGAGTCG-3’ for BCTV C4-G2A; 5’-CACCATGGGGTGCCTCATCTCCATG-3’ and 5’-CTAAATGCTGGCCCTCCCCCCT-3’ for EACMV AC4; 5’-ATGGCGTGCCTCATCTCCATG-3’ and 5’-CTAAATGCTGGCCCTCCCCCCT-3’ for EACMV AC4-G2A.

Briefly, different combinations were transformed into yeast AH109 cells, which were then grown on synthetic defined medium lacking the selected amino acids. Growth of yeast colonies was checked after 72 hours of incubation at 30°C. All experiments were repeated three times

### *Agrobacterium*-mediated viral inoculation

TYLCV infectious clone was described in (Zhao et al., 2021). BCTV infectious clone was purchased from the American Type Culture Collection (ATCC, Manassas, USA). Viral inoculations were performed as described previously (Zhao et al., 2021). Briefly, *Agrobacterium* carrying TYLCV or BCTV infectious clone was syringe-inoculated in the stem of *N. benthamiana* or tomato plants. The inoculated plants were grown in a growth chamber in a 16-h day/8-h dark cycle.

### Elicitor and inhibitor treatments

The flg22 peptide (TRLSSGLKINSAKDDAAGLQIA) (Gómez-Gómez et al., 2019) was dissolved in distilled water at 10 mM as a stock solution. DPI (Sigma, USA), NADPH-oxidase inhibitor, was dissolved in DMSO at 10 mM as a stock solution (Ding et al., 2019). DMTU (Sigma, USA), ROS scavengers dimethylthiourea, was dissolved in distilled water to a 150 mM stock solution (Ding et al., 2019). Brefeldin A (BFA, Invitrogen, USA), an inhibitor of the COPI pathway, was dissolved in DMSO as a 10 mM stock solution (Feng et al., 2016).

Elicitor and inhibitor treatment assay were conducted as described (Ding et al., 2019; Zhai et al., 2021) with some minor modifications. Agroinfiltrated *N. benthamiana* leaves expressing the target protein(s) were treated with flg22 (1 μM), DPI (50 μM), DMTU (5 mM) or BFA (20 μM) at 40 hpai, separately. All mock controls were conducted with 0.2% DMSO or distilled water. The images were taken at 2 hours (DPI and DMTU) or 6 hours (BFA) after elicitor and/or inhibitor treatment. All of the treatments were done independently at least three times.

### Isolation of the chloroplast fraction

Chloroplasts were isolated from *N. benthamiana* plants expressing the protein of interest as described with some minor modifications (Medina-Puche et al., 2020). Briefly, the equal amounts of leaf tissues were incubated with 50 mL chloroplast isolation buffer (50 mM HEPES-KOH pH 8, 5 mM MgCl_2_, 5 mM EDTA pH 8, 5 mM EGTA pH 8, 10 mM NaHCO_3_, 0.33 M D-sorbitol, and Plant Protease Arrest (GBiosciences) at 1:100 dilution). The lysates were transferred to a new tube followed by centrifuging at 400xg for 8 min at 4°C. Then the pellet was resuspended with isolation buffer and added into a 40%/80% Percoll gradient. Intact chloroplasts were carefully collected between tep layers and washed with HS buffer (50 mM HEPES-KOH pH 8, 0.33 M D-sorbitol, and Plant Protease Arrest (GBiosciences) at 1:100 dilution). The chlorophyll concentration was measured as described in (Medina-Puche et al., 2020). The isolated chloroplasts were lysed in HM buffer (10 mM HEPES-KOH pH 8, 5 mM MgCl_2_, and Plant Protease Arrest (GBiosciences) at 1:100 dilution). Total lysates with the same amount of chlorophyll were transferred to new tubes and kept on ice for 10 min. The pellet was washed with HS buffer after centrifuged at 2,600xg for 5 min at 4°C. Then the pellet was resuspended with extraction buffer (20 mM HEPES-KOH pH 7.4, 2 mM EDTA pH 8, 2 mM EGTA pH 8, 25 mM NaF, 1 mM Na_3_VO_4_, 150 mM NaCl, 10% Glycerol, 0.5% TritonX-100, and Plant Protease Arrest (GBiosciences) at 1:100 dilution), and incubated on ice for 30 min. The total proteins were resuspended in the same volume of 1x Laemmli SDS sample buffer. Equal volumes of the solubilized protein fractions were detected by a polyclonal anti-GFP antibody (GenScript, USA). Rubisco large subunit was used as a loading control.

### Immunoprecipitation (IP) using total chloroplast proteins

Monoclonal anti-GFP antibody (Roche, Basel) or polyclonal anti-COPIγ antibody (Agrisera, Sweden) was added to Protein A Sepharose CL-4B (GE HealthCare, USA) and incubated for 2 hours at 4°C to prepare antibody-bound beads. The total proteins extracted from chloroplast fractions as described above were filtered through a 0.22 μm Millipore Express PES membrane, added into the antibody-bound beads, and incubated at 4°C overnight. The beads were washed four times after incubation. Finally, the washed beads were resuspended in the same volume of 1x Laemmli SDS sample buffer. Proteins were detected by a polyclonal anti-COPIγ antibody (Agrisera, Sweden) or a polyclonal anti-GFP antibody (GenScript, USA).

### The plasma membrane (PM) fractionation assays

PM proteins were isolated from *N. benthamiana* cells as described with some minor modifications (Jacob et al, 2021). In brief, the equal amounts of leaf tissues were homogenized with sucrose buffer (20 mM Tris pH 8.0, 0.33 M sucrose, 1 mM EDTA, 5 mM DTT, and Plant Protease Arrest (GBiosciences) at 1:100 dilution) and incubated on ice for 20 min. The lysates were centrifuged at 2,000xg for 5 min at 4°C and the supernatant was labeled as total protein fraction. Then the supernatant was then centrifuged at 16,000xg for 1 h at 4°C, the pellet was resuspended with Buffer B (Minute™ PM protein isolation kit, Invent Biotechnologies, USA) as total membrane fraction. The total membrane fraction was centrifugated at 7,800xg for 5 min at 4°C, and the supernatant was transferred to a new tube and mixed with Phosphate buffered saline (PBS) buffer. The pellet was dissolved with Minute^TM^ Non-Denatured Protein Solubilization Reagent (Invent Biotechnologies, USA) after centrifugation at 16,000xg for 1 h at 4°C and labeled as the PM fraction. Protein concentrations were measured using the Bradford assay (Pierce™ Detergent Compatible Bradford Assay Kit, Thermo Scientific, USA), and equal amount of proteins were loaded on a SDS-PAGE gel for western blot.

Anti-PEPC antibody (Agrisera, Sweden) and anti-ATPase antibody (Agrisera, Sweden) were used as a marker of cytoplasmic or PM protein, respectively. The ratio of C4-YFP/ATPase was set as 1.

### Isolation of tomato protoplast

Protoplasts were isolated from mock- or TYLCV-infected tomato plants following protocols described in (Domozych et al., 2020) with some modifications. Briefly, plant leaves were cut into 0.5 mm strips and digest in the enzyme solution (1.5% cellulase R-10, 0.5% maceroezyme R-10, 0.4 M mannitol, 20 mM KCl, 20 mM MES pH 5.7, 10 mM CaCl_2_). After 3 hours of incubation, an equal volume of W5 solution (154 mM NaCl, 125 mM CaCl_2_, 5 mM KCl, 2 mM MES pH5.7) was added to facilitate protoplast centrifugation. The solution was filtered through a 75 μm nylon mesh into a new tube. Protoplasts were collected by centrifugation at 200xg for 2 min, and the pellet was washed with W5 solution twice. Protoplasts were next resuspend in 0.5 mL W5 solution by gently shaking. Protoplasts were counted under the light microscope by using a hemacytometer.. Protoplasts were kept on ice for 30 min to allow recovery from isolation stress and resuspended in MMg solution (0.4 M mannitol, 15 mM MgCl_2_, 4 mM MES pH5.7) at a density of 10^5^/mL.

### Immunofluorescence staining of COPIγ in tomato protoplasts

Immunofluorescence microscopy was performed as described in (Zhang et al., 2018) with some minor modifications. 15 μL 1% polyethyleneimine was added in a shallow well of microscope slide and incubate for 15 min at RT. Polyethyleneimine solution was removed and 20 μL protoplasts were added in the well and incubated for 20 min. After incubation, IM buffer (1% Non-fat Milk, 0.2% Gelatin, 0.05% BSA, 0.15M NaCl, 0.05M HEPES, 0.5% Tween 20) was added into the well, incubated for 20 min. The well was washed two more times with IM buffer. The anti-COPIγ primary antibody was added to 20 μL IM buffer at 1:100 dilution. The antibody/IM buffer was added into the well and incubated overnight at 4°C. Alexa Fluor 488-conjugated anti-rabbit antibody was added after the wells were gently washed with IM buffer three times, and incubated for 1 h at RT. The well was subsequently washed with IM buffer for three times and sealed with cover slip and a nail polish.

Fluorescence images were observed with a ZEISS LSM 880 laser scanning confocal system. The excitation wavelength for SlCOPIγ signal was 488 nm.

### Quantitative PCR analysis

Total RNA was extracted from tomato or *N. benthamiana* leaves using a hot phenol method (Domozych et al., 2020). RNA samples were treated with M-MuLV Reverse Transcriptase following manufacturer’s instructions (GenScript, USA). The synthesized cDNA was used in qRT-PCR assays to determine transcript levels of the target genes.

The coat protein gene (5’-TATTGTTCGTTGTGTTAGTGATG-3’ and 5’-TGATTAGTGTGATTCTGCTTCT-3’ for TYLCV coat protein; 5’-AACTTCAAATGCTTCGTCACAG-3’ and 5’-TGATATGTTGGGTGCTGGTG-3’ for BCTV coat protein) was detected to represent the accumulated viral genomic DNA from mock-, TYLCV-, or BCTV-inoculated tomato leaves at different time points. The actin gene, *SlActin* (5’-TGTCCCTATCTACGAGGGTTATGC-3’ and 5’-AGTTAAATCACGACCAGCAAGAT-3’), was used as an internal control to test the relative expression levels of viral coat protein gene. SYBR Green supermix (Bio-Rad, USA) was used for qRT-PCR and qPCR. Each experiment was performed in 3 biological replicates and repeated three times. Quantitative PCR were performed with an Applied Biosystems 7500 real-time PCR detection system (ThermoFisher Scietific, USA) and Applied Biosystems 7500 software version 2.0.6 was used for data analysis.

### Quantification and statistical analysis

For western blots, the intensity of each band was analyzed with the ImageJ software (https://imagej.nih.gov/ij/). “Grey Mean Value” of “Region of Interest (ROI)” (contain the specific band of the row) was defined for each blot in chloroplasts lysis or total lysis. The intensity of protein band, cpC4-YFP in chloroplast lysis expressing C4 was set at 1. Values represent the average of three repeated experiments.

All statistical analyses were done with ORIGIN 8 software. Calculated values were from three different leaves of three independent plants. Data are mean ± SD (standard deviation). A one-way ANOVA with Fisher’s least significant difference tests was used to analyze the differences, and *: p-value ≤ 0.05 is to show a statistically significance.

Stromules were quantified from a 20-30 slice z stacks taken by confocal microscopy (Kumar et al., 2018; Caplan et al., 2015). ZEISS LSM browser software was used to generate 2D maximum intensity projections. Photoshop was used to count numbers of stromules and chloroplasts from 30 cells and the ratio of stromules/chloroplasts was calculated. All of the treatments were performed independently at least three times.

To determine if a cell has perinuclear clustering of chloroplasts, perinuclear region was selected and the number of chloroplast was counted in Photoshop. A cell has more than 4 chloroplasts surrounding the nucleus was deemed as positive. In total, 50 cells (from Figure 3C) or 30 cells (from Figures 5B, 5F, 7E) were counted and the percentage of perinuclear-clustering/total cells was shown as “Percentage of nuclei surrounded by >4 chloroplasts”. P values were calculated from three repeats.

The intensity ratio of PM- and chloroplast-localized C4/AC4-YFP in the absence or presence SlCOPIβ (from Figures 3B and 7A) or upon chemical treatment (from Figure 5E) was calculated based on the intensity quantification by ImageJ, as described in (Medina-Puche et al., 2021). Briefly, the same size of “Region of Interest” was set, “Grey Mean Value” from the “Set Measurements” was chosen, and the values were obtained. Three regions of chloroplasts and three regions of the PM were selected in one cell. The average intensity values of the PM and chloroplast were calculated, separately and used to calculate the intensity ratio of PM/chloroplasts. Bars represent SD of n=6 from 6 images. *: p≤ 0.05 is to show a statistically significance between samples based on a one-way ANOVA test.

### Accession numbers

Sequence data from this article can be found in NCBI database under the following accession numbers: SlCOPIβ (XM_026027730), SlCOPIδ (XM_010316171), and SlCOPIε (XM_004231527).

## Acknowledgements

We thank Dr. Rosa Lozano-Duran (Department of Plant Biochemistry, Eberhard Karls University, Tubingen, Germany) for generously providing constructs BCTV-C4-GFP, BCTV-C4-G2A-GFP, EACMV AC4-GFP, and EACMV-AC4-G2A-GFP. We thank Dr. Janet Webster at Virginia Tech, USA, and Dr. Maya Schuldiner at Weizmann Institute of Science, Israel for critically reading the manuscript. We thank everyone in the Zhou and Wang Labs for insightful discussions.

## Author Contributions

W. Z., Y. J., Y. Z., and X. W. designed research; W. Z., Y. J., and X. W. performed research; W.Z., Y. J., Y. Z., and X. W. analyzed data; and W. Z., and X. W. wrote the paper.

